# The influence of pH on the growth and on the formation of nutrient-stress induced scum-forming blooms in cyanobacterial cultures

**DOI:** 10.64898/2026.04.07.716915

**Authors:** Julien Dervaux, Philippe Brunet

## Abstract

The growth of cultures and formation of mucilage blooms in reaction to salt stress of cyanobacterial cultures are investigated with a focus on the influence of pH. In non-buffered medium, cultures show their pH increasing from 6.5 just after inoculation, up to 11 during the exponential phase. We record the time-evolution of concentration and pH, with different initial OD_0_. In a second set of experiments, we extract the doubling time of the unbuffered cultures in comparison with those inoculated in pH-buffered BG11 media at four different pH from 6.3 to 10.5 : in the most acid media, all cultures die or grow very slowly. At pH = 10.5, we obtain the fastest growth for all four strains, allowing to qualify these cyanobacteria as being alkaliphiles, though for all strains with comparable initial OD_0_, the doubling time is shorter for unbuffered cultures. Following a previous study [31]), we finally investigate the influence of pH on mucilage formation and biomass uplift induced by salt stress, involving EPS floculation by cations. Our results show that operating in buffered media significantly influences the mucilage formation, though the observed regimes cannot be simply correlated to the pH value.

## INTRODUCTION

The acidification of freshwater bodies, which mainly originates from the dissolution of CO_2_, NO_*x*_ or SO_*x*_, is less known of the general public than its crucial counterpart in oceans [1]. Though, the decrease of pH can be harmful for many freshwater species, and can threaten their habitats. On the other hand, aquatic plants and photosynthetic microorganisms are able to mitigate this acidification by assimilating CO_2_, either by photosynthesis or by mineralisation, and thus constitute a powerful sink of carbon. Hence in certain conditions, some microbes like cyanobacteria can benefit from the rise of atmospheric CO_2_ concentration [2–4].

In this sense, the growth of these photosynthetic microorganisms, of either non-axenic or axenic cultures, depends on the concentration of inorganic carbon, which can be assimilated by cyanobacteria or micro-algae either as dissolved CO_2_, forming carbonic acid H_2_CO_3_, or in ionic forms as bicarbonate 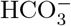 and carbonate 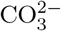. This makes pH a factor of utmost importance for the survival and growth of these micro-organisms [5]. Within media of pH under 6.5, many cyanobacterial cultures are generally unable to grow or even to survive. On the other hand, cyanobacteria can endure extreme media [6], and are even able to modify their ambient environment [7– 10], presumably for their own advantage. This modification should be especially significant in freshwaters, since they often offers stagnant conditions, with weak swirling and sometimes confined spaces. How such microorganisms can adapt to different acidic or alcalin conditions, is fundamental not only for their survival but also for the resilience of many ecosystems [8, 11, 12], as well as for more applied prospectives, eg. the optimisation of culture conditions in photobioreactors [6]. Though, investigations on the growth of cyanobacteria, microalgae or any other photosynthetic microbes under specific pH conditions, remain scarce [13–16].

The propension for photosynthetic microbes to modify their environment is of course much dependent on the photosynthetic activity, being itself dependent on the eutrophication. Indeed, a relatively poor medium, with low microbe concentration, keeps a CO_2_ level close to that at equilibrium with ambient air, since its consumption rate is much smaller than its rate of supply from air. Therefore, the CO_2_ can be readily transformed to carbonic acid H_2_CO_3_ and the value of pH remains around or inferior to 7. Conversely, in eutrophic media the CO_2_ can be readily depleted, especially because in water, the diffusion of CO_2_ is much slower than in air. Hence it has been measured that the value of pH of the surrounding water could rise up to 10 or even beyond [9, 10].

One of the most direct and straightforward way to quantify the influence of pH is to measure the growth-rate of a culture under fixed pH. These measurements are generally carried out with axenic cultures in well-controlled laboratory conditions. Then, one can wonder how results obtained within such idealised situations can be relevant or insightful for environmentally-oriented purposes. Indeed in natural habitats, microbes generally evolve in ambient conditions that can significantly fluctuate in space and time. Therefore, the adaptation to changing conditions should be a key for their survival. Amongst the different ways cyanobacteria could adapt, their ability to modify the ambient pH should be a powerful asset. Not only the drift of pH toward an optimal value could enables a more efficient assimilation of inorganic forms of carbon [10], but also a medium becoming alcaline has been shown to offer a significant advantage to cyanobacteria over other micro-organisms [14]. In an applied prospective, such alcaline conditions are especially interesting to mitigate contaminations in photo-bioreactors. Furthermore, alcaline conditions are susceptible to improve the ability of cyanobacteria to produce siderophores [6, 17].

But the question of the very occurrence of such a pH modification by the cyanobacteria themselves remains to be quantitatively evidenced in details [9]. Furthermore in terms of a quantitative study, to measure the time-evolution of pH of an unbuffered culture would constitute an alternative way to evaluate such an adaptation. And beyond providing experimental evidences of such a drift of pH over the time of the culture growth, a very important point is to identify the underlying mechanisms of pH modification.

In this prospect, it is important to remind that under low CO_2_ and high pH conditions, many photosynthetic micro-organisms (PMOs), and cyanobacteria in particular, ensure the uptake of inorganic carbon using a Carbon Concentration Mechanism (CCM) [7, 18–27]. To circumvent the limitations of a low CO_2_ concentration, the CCM ensures that RuBisCO is surrounded by CO_2_ instead of O_2_ in the carboxysomes, in order to prevent the oxygenase reaction. More precisely, the CCM enables to access the pool of 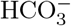 to achieve photosynthesis, the bicarbonate becoming CO_2_ within the cells. The reaction lead to the exportation of OH^−^ outside the cells [19–23, 26, 28, 29], in turn inducing the increase of external pH. Since the form 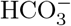 is promoted by alcaline conditions, relatively to H_2_CO_3_, the CCM could be one of the key ingredient for a pH regulation toward optimal conditions for the growth of cyanobacteria. Recently, it has even been postulated that pH could play a crucial role in the energetic efficiency of CCM [5]

Despite very recent studies at the cell scale [30], various questions remain unanswered : how does the pH evolve with time for different culture strains and initial concentration ? Does pH reach an equilibrium value or does it endlessly drift until the senescence of the culture ? Does this final value correspond to an optimum for the growth of the culture ? How does the culture react to a sudden change of pH ? How does pH influence the appearance of scum-forming blooms induced by salt stress reported in previous experiments [31–37] ?

In this study, we present results of the influence of pH on the growth of cultures for four axenic strains of bloom-forming, freshwater cyanobacteria. The four strains are *Microcystis Aeruginosa* PCC 7005, *Synechocystis sp*. PCC 6803, *Anabaena sp*. PCC 7120 and *Synechococcus Elongatus* PCC 7942. The choice for these specific strains are motivated by that they form blooms in the environnement. Also, we opted for strains encompassing different morphologies, from round to elongated or filamentous. Table 1 summarises the references of the used strains. For each strains, and different initial optical densities (OD), we measured the concentration and pH of the different cultures during up to 4 weeks after inoculation, every 2 or 3 days.

**TABLE I:** Name, references and characteristics of the axenic strains in this study.

| Name | Strain ref. | Shape | Observations |
| --- | --- | --- | --- |
| *Microcystis Aeruginosa* | PCC 7005 | Colonial | With vacuoles |
| *Synechocystis sp.* | PCC 6803 | Unicellular spheroid | - |
| *Anabaena sp.* | PCC 7120 | Filamentous | Nitrogen fixing |
| *Synechococcus Elongatus* | PCC 7942 | Unicellular ellipsoid | Nitrogen fixing |

The paper is organised as follows : after describing the experimental methods, we present results of the cyanobacterial concentration and pH for various strains at different initial concentration, versus time, in unbuffered medium. All cases generally show an exponential growth, from which a doubling time is extracted. Then, we present the concentration versus time, of the same strains inoculated in pH-buffered medium. We also measure the time-evolution of pH of cultures under various initial conditions after a sudden acidification, together with the time-evolution of their OD. In a more qualitative way, we show that pH has an important influence on the appearance of scum-forming blooms. We finally discuss on the possible consequences of these findings in environmental and practical situations.

## MATERIALS AND METHODS

### Strains and culture conditions

Cultures were growth in unbuffered or pH-buffered medium BG11 broth and Trace Metals solution, under a 14h/10h light cycle at 13.5 *µ*E.m^−2^.s^−1^ at 25°C. The composition of the BG11 medium (1X) is as follow : CaCl_2_.2H_2_O (36.7mg/L), citric acid (5.6 mg/L), K_2_HPO_4_ (31.4mg/L), Na_2_Mg.EDTA (1mg/L), C_6_H_8_FeNO_7_ (6mg/L), MgSO_4_ (36mg/L), Na_2_CO_3_ (20mg/L), NaNO_3_ (1500mg/L). The trace metal solution is composed of : H_3_BO_3_ (2860 mg/L), MnCl_2_4H_2_O (1810 mg/L), ZnSO_4_7H_2_O (222 mg/L), Na_2_MoO_4_2H_2_O (390 mg/L), CuSO_4_5H_2_O (79 mg/L), Co(NO_3_)_2_6H_2_O (49 mg/L). The unbuffered medium has a pH of 7.05 ± 0.05 just after inoculation. Some characteristics of the four strains used here, are indicated in Table I.

Buffers were chosen in order to be biocompatible, according to the criteria of Good *et al*. [38]. We opted for four bio-buffers (acquired from Sigma Aldrich), covering a range of pH from 6.2 to 10.4, and denoted thereafter as MES, MOPS, TAPS and CAPS (sorted by increasing pH). Their complete names and properties are listed in Table II.

**TABLE II:** Name and main characteristics of pH bio-buffers.

| Name | Abbreviation | pKa (at 25°C) | Ambient pH |
| --- | --- | --- | --- |
| 2-(N-morpholino)ethanesulfonic acid | MES | 6.15 | 6.34 |
| 3-(N-morpholino)propanesulfonic acid | MOPS | 7.20 | 7.40 |
| (tris(hydroxymethyl)methylamino)propanesulfonic acid | TAPS | 8.44 | 9.50 |
| N-cyclohexyl-3-aminopropanesulfonic acid | CAPS | 10.40 | 10.42 |

The buffered media are prepared at 25 mM dilution of each acid buffer, completed with 1M NaOH drop-by-drop until the pH stays constant at ± 0.1 after the addition of 2 consecutive drops. In practice, the medium pH reaches a value roughly corresponding to the pKa at 25°C, although with a slight shift, see Table II. This shift is particularly large for the TAPS buffer with a final pH=9.5 for a pKa = 8.44. We systematically check that the pH values remain unchanged after the inoculation of the cultures.

### Biomass measurements

Optical density measurements were carried out using a Jenway 7310 spectrophotometer, putting liquid cultures samples in a cuvette with a 1 cm light path, at optical wavelength 580 nm [39, 40]. Cell counting was performed with a Malassez cell on an inverted microscope and gave a concentration of 2 × 10^7^ cell/ml at an optical density OD = 1. Using an average cell diameter of 5 *µ*m, this corresponds to a volume fraction of roughly 10^−3^. Since the relationship between OD and cyanobacteria concentration is no longer linear if OD*>* 1, we systematically re-dilute the sample in fresh medium before doing the measurement, so that OD of the diluted sample remains inferior to 1.

### pH measurements

All measurements of pH followed the same protocol. First, the probe of a pH-meter (Hanna, probe Hi1131B) was abundantly washed with micro-pure water, then dried with clean room wiper and plunged into the erlenmeyer containing the culture until the pH keeps a stable value, generally after less than 30 s. The culture is slightly shaken by hand, and installed below the flame of a Bunsen burner to prevent any contamination. The pH-meter was frequently calibrated with buffers at pH = 4, 7 and 10.5.

Except for measurements quantifying the influence of the time in the day on pH, all measurements were carried out between 7 PM and 8 PM.

## EXPERIMENTAL RESULTS

### Cultures growth and pH time-evolution in a non-buffered medium

We first measured the time-evolution of the cell concentration, quantified by the optical density at 580 nm (denoted OD), with the different strains growth a nonbuffered medium. We chose different initial cell concentration OD_0_, to quantify the influence of the initial conditions. The typical overall duration of a measurement campaign was between 22 and 28 days after inoculation, or roughly 500 to 700 hours.

Panels (a,b,c) of Fig.1 show the time-evolution of these optical densities, represented in lin-log plots. In most situations, the optical density shows an exponential growth starting just after inoculation. Sometimes, a lag phase of 1 to 2 days in duration is observed before entering the exponential growth, especially for cultures with low initial concentration. The saturation phase occurs roughly after 400 hours (or ∼ 16 days). For each culture, we extracted the doubling time *τ*_2_ within the exponential phase. We present these doubling times *τ*_2_ as a function of the initial OD_0_ in panel (a) of Fig. 2. We note that the doubling time increases monotonically with the initial cell concentration.

**FIG. 1:**
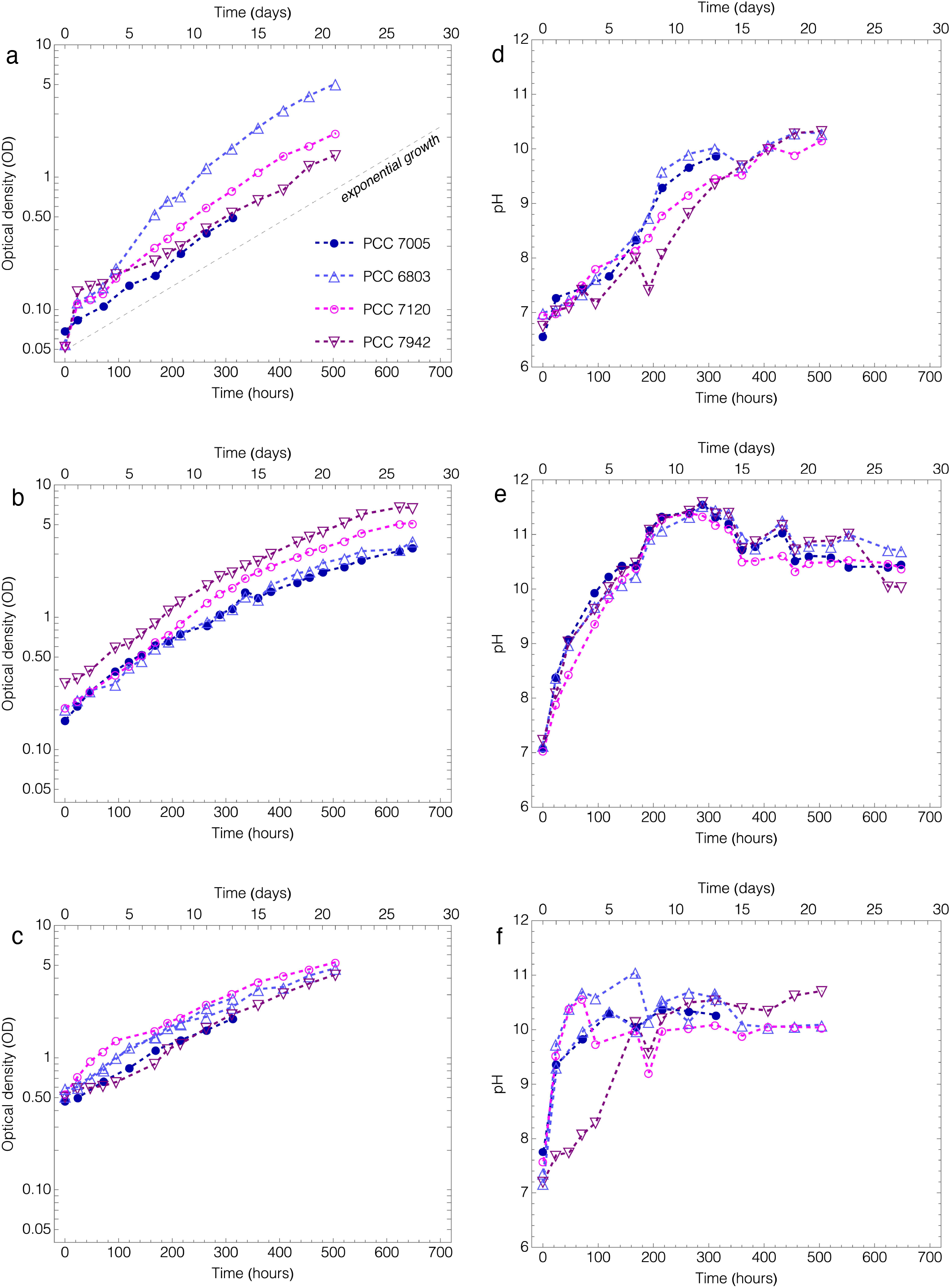
(a,b,c): concentration of cultures, in optical density at 580 nm (OD), versus time after inoculation, in lin-log axes, of four different strains grown in unbuffered BG11. (a) Initial OD around 0.05. (b) Initial OD between 0.2 and 0.3. (c) Initial OD between 0.5 and 0.6. All series show a classical exponential growth phase, as suggested by the dashed line in panel (a). (d,e,f): pH of cultures versus time after inoculation, of four different strains grown in unbuffered BG11. The data series (d,e,f) correspond to the OD shown respectively in (a,b,c).

**FIG. 2:**
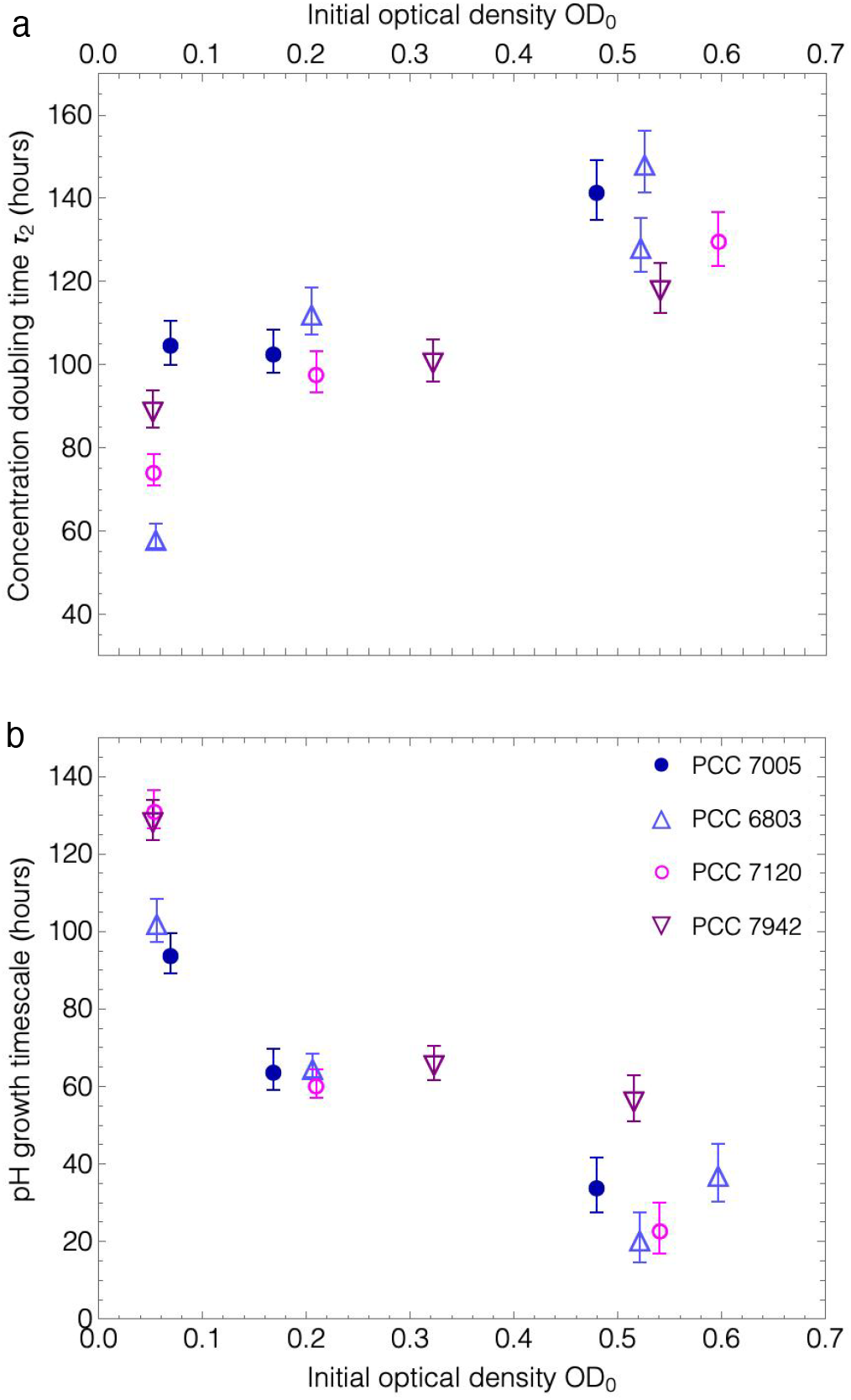
(a): concentration doubling time of unbuffered cultures versus initial optical density OD_0_. (b): timescale of growth of the pH extracted during the initial phase, prior to saturation, versus initial optical density OD_0_ for the four cyanobacterial strains.

Panels (d,e,f) of Fig.1 show the time evolution of the pH of the cultures corresponding to the series of data shown in panels (a,b,c) of Fig.1: panel (d) shows data for an initial OD around 0.05, panel (e) shows data for an initial OD between 0.2 and 0.3 and subfigure (f) shows data for an initial OD between 0.5 and 0.6. For all measured cultures, the initial pH takes values ranging between 6.8 and 7.2, and then increases with time in the very first days. Regardless of the cyanobacterial strain or the initial concentration OD_0_, the pH reaches a saturation value - ranging between 10 and 11, within a timescale of 5 to 20 days after inoculation. Let us remark that the value of OD_0_ influences the sharpness of the initial increase. Panel (b) of Fig. 2 shows the timescale of pH growth (i.e the time needed for the pH to increase by 1 unit) versus OD_0_, extracted from linear fits within the growing phase (before saturation) in panels (d,e,f) of Fig.1. The suggested trend is that the initial growing phase of pH is sharper in time for higher initial concentration, regardless the strain.

Let us mention that the values of pH in all the previous plots, including in the saturation stage, are all measured between 7 PM and 8 PM. Since the cultures are subjected to a daily cycle of light/darkness of 14h/10h - with lights starting at 7 AM, the cultures have then been exposed to light for 12 to 13 consecutive hours of light and hence cyanobacteria make photosynthesis and consume CO_2_ and HCO_3_-, leading to the increase of pH. Conversely during the darkness phase, respiration should induce the decrease of pH. In this sense, we check that the time of the day indeed influences the value of the pH in a significant way. Fig.3 shows typical example of this daily variations for four cultures with different strains, once the pH has saturated (i.e ∼ 2 weeks after inoculation). Depending on the strains, the daily pH variations range between 0.8 and 1.2.

**FIG. 3:**
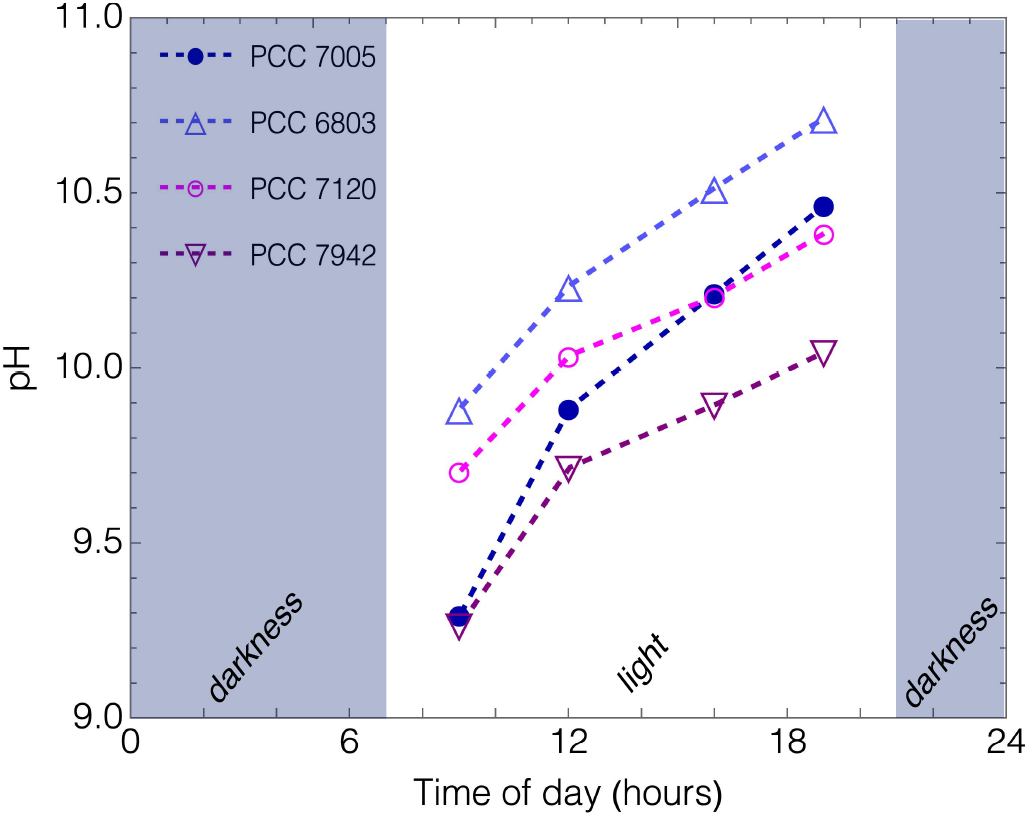
Daily variation of pH of four cultures of the four different strains at 9am, 12pm, 16pm and 19pm during the same day. Light is turned on at 7am and turned off at 21pm.

### Cultures growth in buffered media

We now present the growth of the four different strains in pH-buffered media. As previously, these measurements were carried out in a long enough period of time, during 33 to 40 days, so that the saturation phase after the exponential growth is observed. Again, measurements were performed between 7 and 8 PM. Initial concentration ranges between OD_0_ = 0.15 and 0.45. This choice of initial concentration is motivated by the fact that, in this range, the doubling times *τ*_2_ in unbuffered medium are roughly the same for all four strains studied here, as can be seen in Fig.2-a. As shown in Fig.4, for the all four strains used, the value of pH has a strong influence on the growth of the culture. We then quantify the growth by extracting the doubling time *τ*_2_ during the exponential phase. Before presenting these quantitative results in Fig.5, let us make a few qualitative remarks on these experiments:

**FIG. 4:**
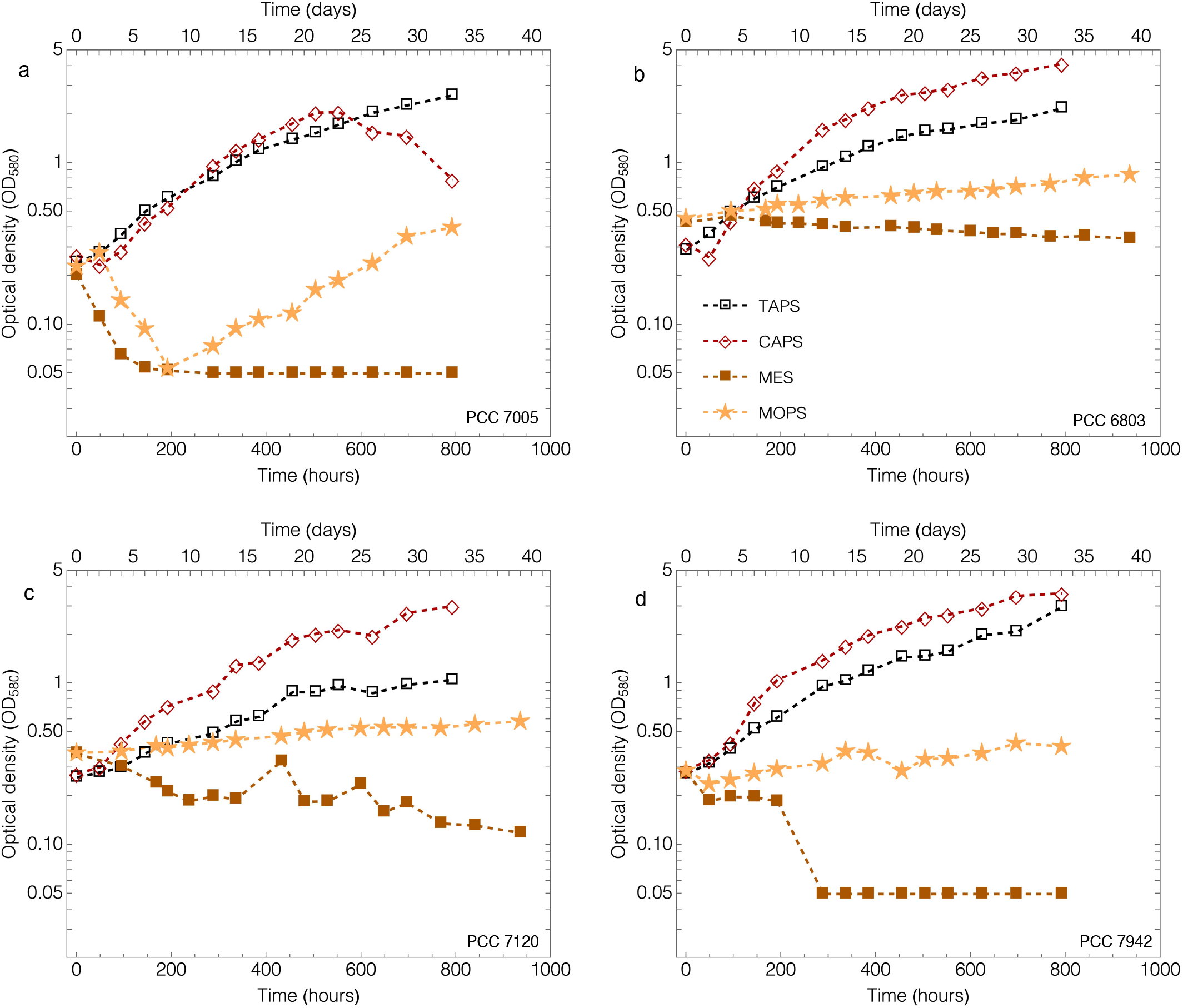
Concentration of cultures, given in optical density (OD) at 580 nm, versus time after inoculation, in lin-log axes, for four different cultures growth in BG11 buffered with MES (pH=6.34), MOPS (pH=7.4), TAPS (pH=9.5) and CAPS (pH=10.42). (a) *Microcystis Aeruginosa* PCC 7005, (b) *Synechocystis sp*. PCC 6803, (c) *Anabaena sp*. PCC 7120 and (d) *Synechococcus Elongatus* PCC 7942. Initial OD ranges between 0.2 and 0.45.

**FIG. 5:**
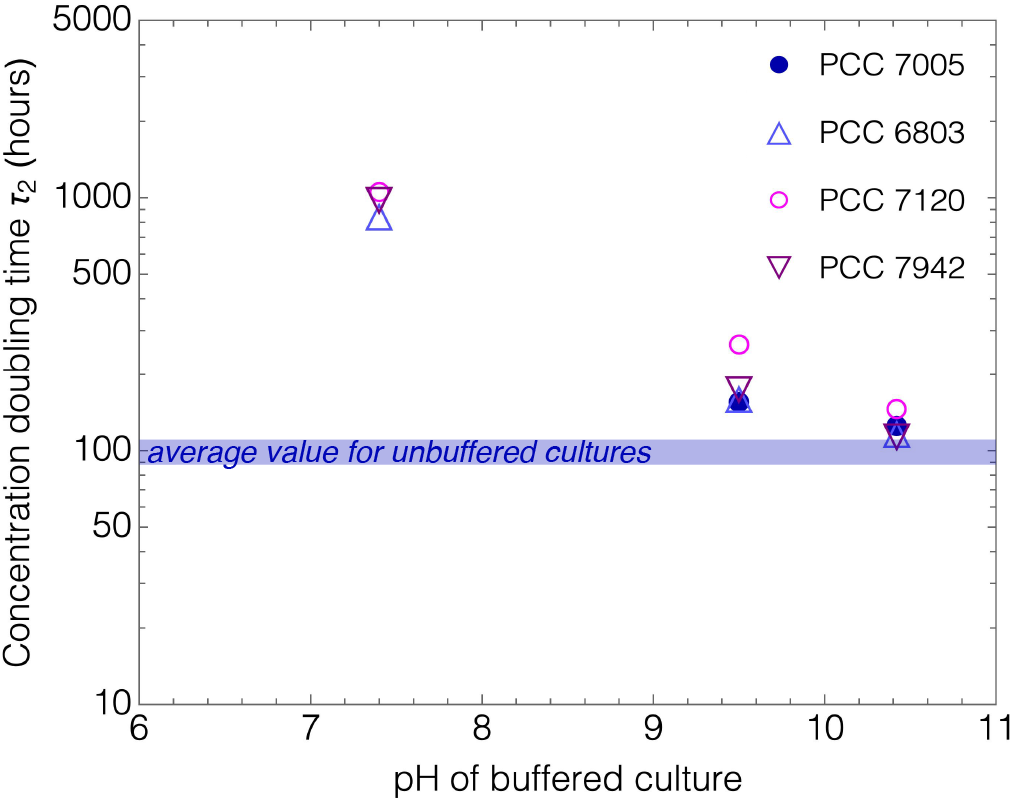
*Top* - Doubling time of the different cultures versus pH. The different symbols stand for different cultures in buffered media, while the grey rectangles represent the ranges of values extracted from growth in unbuffered media: dark grey one encompasses cultures with intermediate values of OD_0_ and the light grey one encompasses values for all OD_0_. *Bottom* - Doubling time of unbuffered cultures versus OD_0_ - the dashed-line circle surrounding those corresponding to the dark grey rectangle.

- All cultures growth in MES-buffered medium (pH = 6.34) showed their OD decaying in time: PCC 7005 (Fig.4-a) and PCC 7942 (Fig.4-d) cultures died after a few days, while PCC 6803 (Fig.4-b) and PCC 7120 (Fig.4-c) progressively declined over the course of the experiment, with an OD decreasing by a factor of 1.5 to 3 compared to its initial value.
- The cultures in MOPS-buffered medium (pH = 7.4) showed a very slow growth during the whole measurement campaign the optical density merely increasing by a factor 1.2 − 1.6, depending on the strains, in more than three weeks. As shown in Fig.5, this leads to very large doubling time approaching ∼ 1000 hours.
- The cultures in TAPS-buffered medium (pH = 9.5) showed a steady growth from the very first days after inoculation.
- The cultures in CAPS-buffered medium (pH = 10.42) showed the fastest growth of all buffers.

Interestingly, if we compare the values of *τ*_2_ in buffered media with those extracted from unbuffered cultures (for comparable initial OD_0_), the later are systematically shorter for all strains used although the *τ*_2_ for CAPS-buffered cultures are comparable in magnitude with those of unbuffered cultures. Fig.5 shows the doubling times versus pH, with the blue band showing the range of values for unbuffered cultures (∼ 100 hours).

Hence, at least for the strains tested here, our cyanobacterial cultures grow faster in unbuffered media than in any buffered ones, including in that at pH = 10.5, close to the plateau value reached in unbuffered media. Although the CAPS-buffered media at pH = 10.5 would correspond to a nearly optimal condition for growth, the growth is faster in a medium with pH freely evolving from roughly 7 to 11.5. Though we cannot propose any precise explanation at this stage, it is important to note that in a buffered medium, the pH remains fixed all day long, including during the darkness phase, while its value drops by 0.8 to 1.2 units in an unbuffered medium, as shown in Fig.3. Therefore, while in a high-pH buffer like CAPS, the growth conditions should be suitable during the photosynthesis phase, these conditions may not be optimal for the respiration phase, since during the latter the production of CO_2_ would rather benefit from less alcaline conditions, see eg. [5].

### The pH-adaptation of a culture after a prompt re-acidification

In order to test the robustness of the alkalisation in cyanobacterial cultures, we check the response of cultures under various initial conditions (time after inoculation, OD, strain) to a sudden re-acidification. From cultures of 3 to 9 weeks after inoculation, corresponding to the exponential growth phase or a later phase, we take a sample of 30 to 40 mL of cultures with OD ranging typically between 3 and 6.5, in which we add 5 to 30 *µ*L of concentrated acetic acid. Before acidification, initial values of pH range between 9 to 11. After acidification, the pH values range from 4 to 7. The experimental conditions for the different experiments are summarized in Table III.

**TABLE III:**
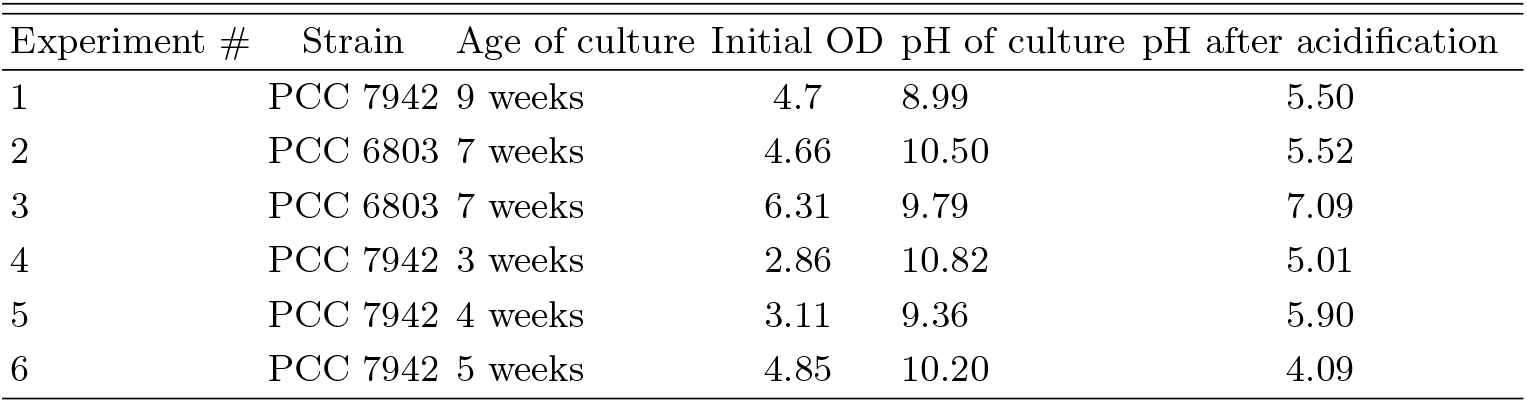
Experimental conditions of the prompt reacidification measurements.

The pH is measured every one to three days between 7 and 8 PM. The OD of each sample is also measured. The results are depicted in figures 6-(a,b). In all tested conditions when the pH was initially decreased to a value around 5 or higher, the pH increases and reaches a final value between 8.5 and 9.5 (not necessarily correlated to its value before acidification) after a duration of roughly 300 to 500 hours, sometimes after an earlier overshoot at up to 11. In all of these cultures except one (where OD drops from 3 to less than 0.5), the OD remains at a relatively stable value with only a slight decrease and even an increase during later stages (*t >* 400 hours). It is only if the pH has been decreased down to 4 that the value of pH does not increase and remains close to 4. Surprisingly, the OD of the tested culture remains constant, only dropping from 4.7 to roughly 3.8 followed by a slight increase later (*t >* 200 hours).

**FIG. 6:**
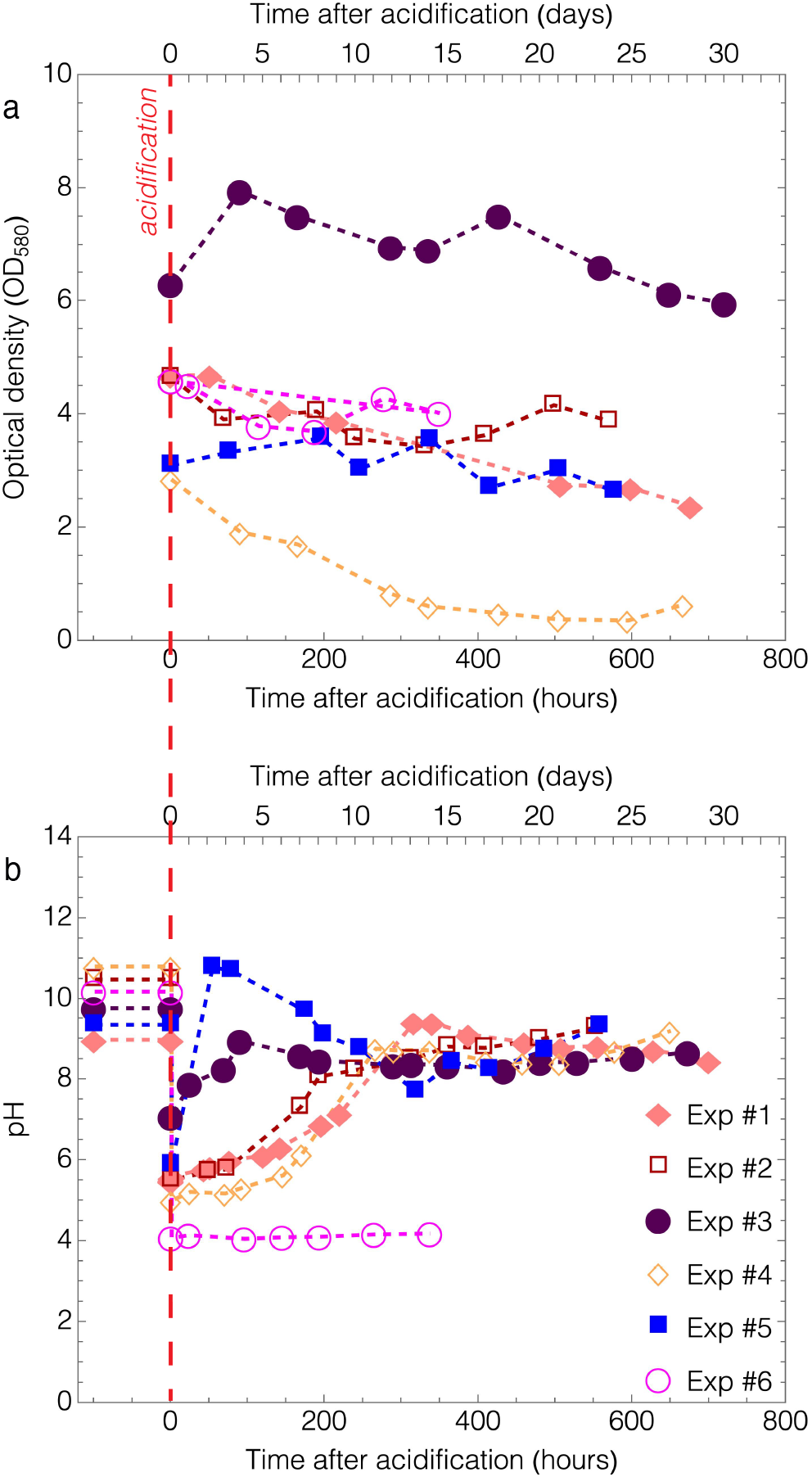
*Top* - Time-evolution of the pH of various cultures after re-acidification (see table III for the conditions). *Bottom* - Corresponding time-evolution of the OD of the cultures.

We conclude that cyanobacterial cultures in their exponential growth phase or in later stage have the ability to counteract a sudden acidification of their medium, by increasing the pH to the alkaline range and sometimes up to a value higher than that prior to acidification. This alkalination is not observed if the pH is decreased down to roughly 4.

### The effect of pH on the formation of mucilage blooms

In a previous study, we showed that the addition of mono- or divalent cations (contained in practice in the BG11 medium) were susceptible to cause the formation of scums in cultures of bloom-forming cyanobacteria *Microcystis Aeruginosa* PCC 7005 and *Synechocystis sp*. PCC 6803 [31]. While axenic strains in laboratory are less prevalent to form colonial structures, this scum was explained by the chelation of unbound EPS with cations, hence ensuring the mechanical stability of the scum. In turn, the continuous production of O_2_ by photosynthesis induced the global rise of the biomass to the top of the surface, since although trapped in their EPS matrix, cyanobacteria keep their metabolic activity. This irreversible bloom is reminiscent of scums observed in the environment, and has been reproduced in various other studies [32–37]

Here, we investigate how the formation of these irreversible scums, and their rise on the top of the water column, could be influenced by the value of pH. Therefore, we repeated experiments in the spirit of [31] in the various buffered media. Though the results are much less quantitative than in our aforementioned study, they provide interesting clues on the role of pH. They also reveal that the formation and the stability of the scum are very different from one cyanobacterial strain to another, so that we opted to present the observations separately for each strain. Some of the strains are more prone to form stable scum with the addition of CaCl_2_ instead of BG11 (eg. PCC 6803 [31]).

The experimental protocol is inspired from that followed in [31] : a sample of cyanobacterial culture is put in a sterilised glass tube and completed to 5 mL with either 100X concentrated BG11 broth or CaCl_2_ at *c* = 1 mol/L. The high concentrations in BG11 and CaCl_2_ enable to add a small enough volume of concentrate (typically from 50 to 500 *µ*L), so that the subsequent drop of culture OD is not significant. The mother culture is chosen to be close to the end of the exponential growth phase, typically 2 or 3 weeks after initial inoculation. The choice is motivated by that the OD has to be high enough (OD between roughly 2 and 3.5 before dilution), and by that older cultures have shown to have a lesser amount of EPS and hence do not build a robust enough EPS matrix [31]. The buffered cultures are prepared by diluting 3.5 to 5 times the concentrated mother culture in buffered media. We check that the pH of the cultures is indeed that of the buffered media.

The initial OD ranges between 0.5 and 2, though in most experiments (especially for buffered cultures) the OD is between 0.5 and 1. Each tube then contains a sample of culture with BG11 or CaCl_2_ at given concentration. Like in [31], we indicate the total concentration of BG11 with respect to the nominal value, from 1X (one tube is always taken as a control) to 10X if the concentration is 10 times higher (the maximal value imposed here), whereas the added concentration in CaCl_2_ is given in mM (the BG11 itself provides CaCl_2_ at 0.23 mM).

Each sample is gently mixed by inverted each tube upside-down three times. The tubes are then placed in an enclosure made of opaque walls to shield them from external perturbations or unwanted light, and subjected to continuous white light at 1000 ±50 Lux (unless otherwise specified), which was found to be in the medium range of intensity for which the rise of biomass was observed. A camera installed in the enclosure takes pictures every 1 to 3 minutes, to generate a time-lapse sequence over a duration of 12 to 24 hours. A typical result is depicted in Figure 7.

**FIG. 7:**
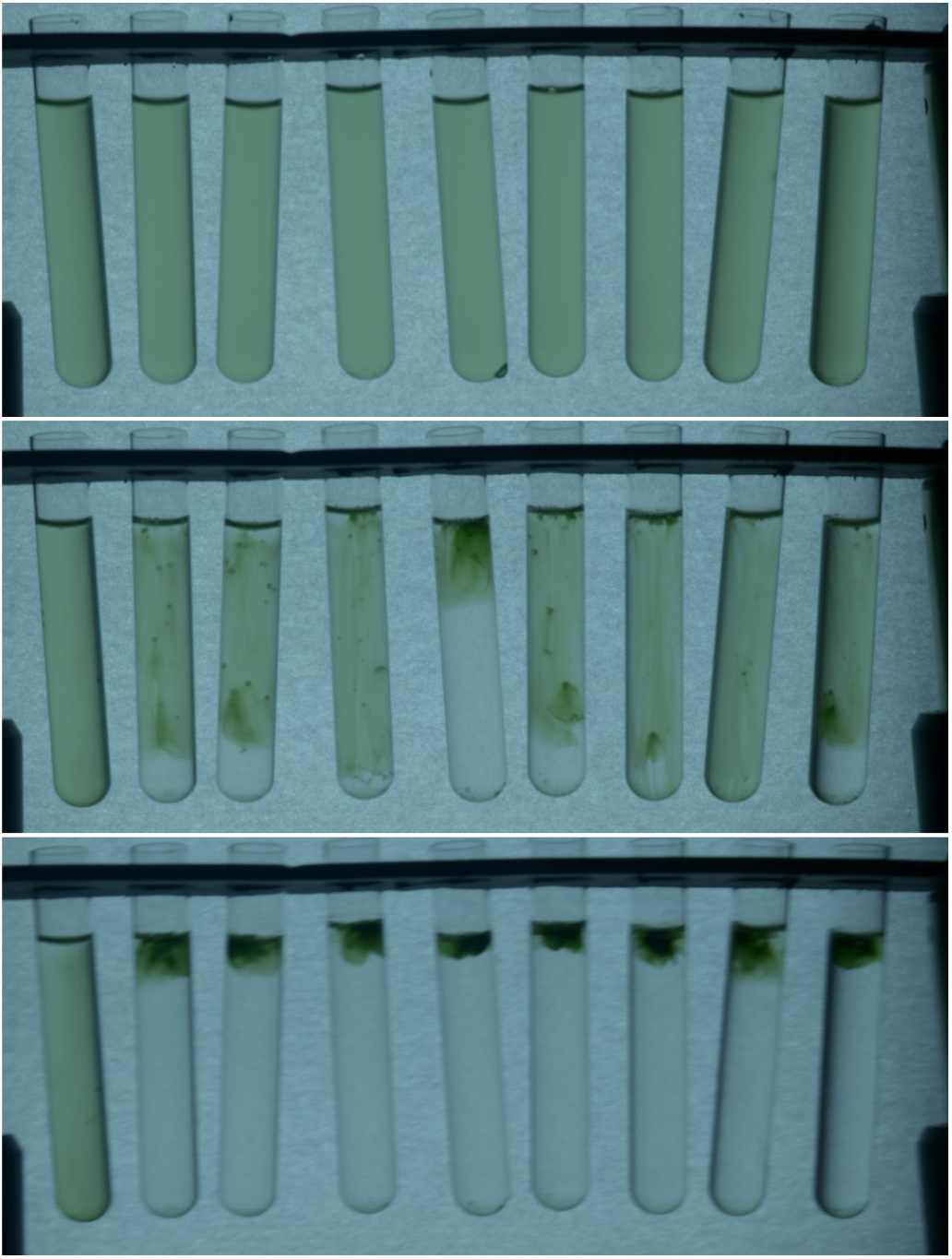
Samples of cultures of PCC 6803 in unbuffered medium, initial OD = 1.35 and their evolution after addition of CaCl_2_ at various concentration, from left to right 0 mM (control tube), 0.5 mM, 1 mM, 1.5 mM, 2 mM, 2.5 mM, 3 mM, 5 mM and 10 mM. From top to bottom : 5 mn, 16 h and 22h after salt addition, showing the formation of O_2_ bubbles lifting the whole biomass up to the surface (IRMB regime), except for the control tube.

In the following, we denote several distinct behavior following the addition of concentrated salts as : irreversible rising mucilage bloom (IRMB) when as observed in [31] the biomass rises to the surface embedded in an EPS matrix with entrapped bubbles, partial (PS) or complete sedimentation (CS) when the biomass sinks, intermittent rise and sink of biomass (IRS) or homogeneous biomass concentration (HB) the latter meaning that almost no visual change is noticed over the course of the recording. We provide videos of the most remarkable observations as Supplementary Information.

#### Microcystis Aeruginosa PCC 7005

We first check the threshold in BG11 for IRMB to be consistent with that measured in [31] for PCC 7005, where it was found the threshold (roughly 4X) depended little on light intensity and initial OD. Several samples of a culture at OD = 0.605 from a mother culture of OD = 2.28 with a dilution ratio of 3.8, are poured in tubes where the BG11 concentration is varied from 1 to 10 X by steps of 1X. The duration of the rise ranges between 60 and 180 mn, depending on the BG11 concentration.

When a similar mother culture is diluted in buffered media, this behavior changes qualitatively. This means that the IRMB regime can be prevented, and replaced by other behavior. Conversely, under some buffered conditions, the IRMB seems to be favoured. All PCC 7005 cultures have initial OD around 0.6 ± 0.05.

In MES buffer, a PCC 7005 culture is found to exhibit the IRMB regime for any concentration in BG11, including in the control tube (1X, no added BG11). Though, the duration of the rise ranges roughly between 60 and 300 mn, but with no obvious dependence on added BG11.

In MOPS buffer, a PCC 7005 culture exhibits partial (PS) or complete (CS) sedimentation. No IRMB is observed. The biomass shows clusters similar to those in IRMB, but they slowly sink instead of rising. Still, bubbles are observed, especially stuck to the walls, but the EPS matrix seems too fragile to retain rising bubbles in between.

In TAPS buffer, a PCC 7005 culture shows a mixed behavior between IRMB, PS and CS, see figure 8. At low BG11 concentration, including in the control tube observed separately, the culture remains homogeneous (HB). The clustering of the biomass caused by the addition of salts, observed above a concentration of BG11 roughly equal to 3.5X. The main cluster either sinks (PS, CS) or rises (IRMB), but with no obvious dependence with salt concentration. Let us mention that, although seemingly homogeneous in Fig. 8, the tube 6X (last in the right) shows CS. Indeed once a slight perturbation is brought, the biomass sinks in a few minutes. The same behavior is observed at 8X and 10X.

**FIG. 8:**
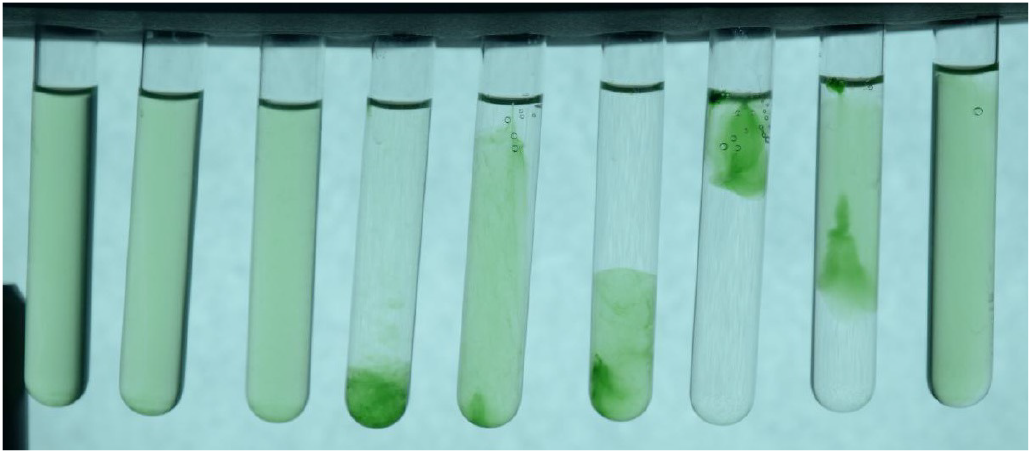
Samples of cultures of PCC 7005 in TAPS-buffered medium, initial OD = 0.6 and their evolution after addition of BG11 at various concentration, from left to right 2X, 2.5X, 3X, 3.5X, 4X, 4.5X, 5X, 5.5X, 6X. The control tube (1X) and those at higher BG11 concentration are observed separately under similar conditions. At 3X or below, the culture remains homogeneous. Above 3X, the cultures show PS, CS or IRMB.

In CAPS buffer, a PCC 7005 culture shows PS and CS behavior. Instead of a single large cluster, the addition of salt induces many milimeter-sized clusters, which average size seems to increase from 2X to 4.5X, leading to a faster and more complete sedimentation at 4.5X. At 5X to 10X, a larger unique cluster forms, but cannot retain bubbles in it. Therefore, similar to cultures at high BG11 concentration with TAPS, the biomass slowly sinks (PS). The control tube remains homogeneous. A video is provided in S.I. for concentrations from 2X to 6X.

#### Synechocystis sp. PCC 6803

Previous experiments revealed that PCC 6803 cultures would not show IRMB after addition of BG11, even at high concentration (up to 50X), but instead would show IRMB after the addition of CaCl_2_ at concentration as low as 5 mM [31]. Therefore, we rather opted to study the influence of pH-buffered media in this latter situation.

First, we check that the observations with BG11 addition would change or not in buffered media. We used a culture with OD = 0.9. In MES and MOPS, the cultures from 1X to 5X do not show visible biomass migration, with only tiny clusters floating below the surface for 5X concentration. In CAPS, these tiny clusters are also observed at 5X, but they rise up the surface, leading to a sort of partial IRMB. In TAPS, a global rise of biomass is observed in specific conditions : from 2X to 5X, an uplifting leading to IRMB is observed, but its duration is much longer than what observed for PCC 7005 : after one day, the uplifting was not yet done and in some tubes, the position of the lower biomass front was somehow frozen. Still, the fastest rise was found for 4.5X while it was slower at higher concentration. For 6X and above, the culture stays homogeneous, as for the control tube.

We then investigate the effect of addition of CaCl_2_. Figure 7 (+ Video in S.I.) shows that all unbuffered cultures, except the control tube, exhibit IRMB. It is to be mentioned that the threshold is as low as 0.5 mM. In details, the biomass cluster appears in a filamentous, slender shape in MES and MOPS. In TAPS and CAPS, the agglomeration appears in a few seconds after addition of CaCl_2_ if the concentration is high enough, several minutes to one hour for a lesser concentration. As a result, the whole biomass sinks before the oxygen bubbles can grow and lift it up. Therefore, a first stage of CS is followed by IRMB. See Figure 9 and videos in S.I. for more details. Concerning the CaCl_2_ concentration threshold, in MES buffer, the IRMB regime is hindered at low to moderate concentration in CaCl_2_. It is only observed above roughly 80 mM, but not clearly as it seems no bubble can grow in the culture. In MOPS, the threshold is measured between 50 and 60 mM (see 9-(Up)) but not in a clear way. In TAPS and CAPS buffers, the IRMB appears at 0.5 mM or above like in unbuffered medium, hence comparable to unbuffered medium. Therefore, not only the buffers influence the way EPS aggregate the biomass, but also the value of pH seems to influence the growth of oxygen bubbles.

**FIG. 9:**
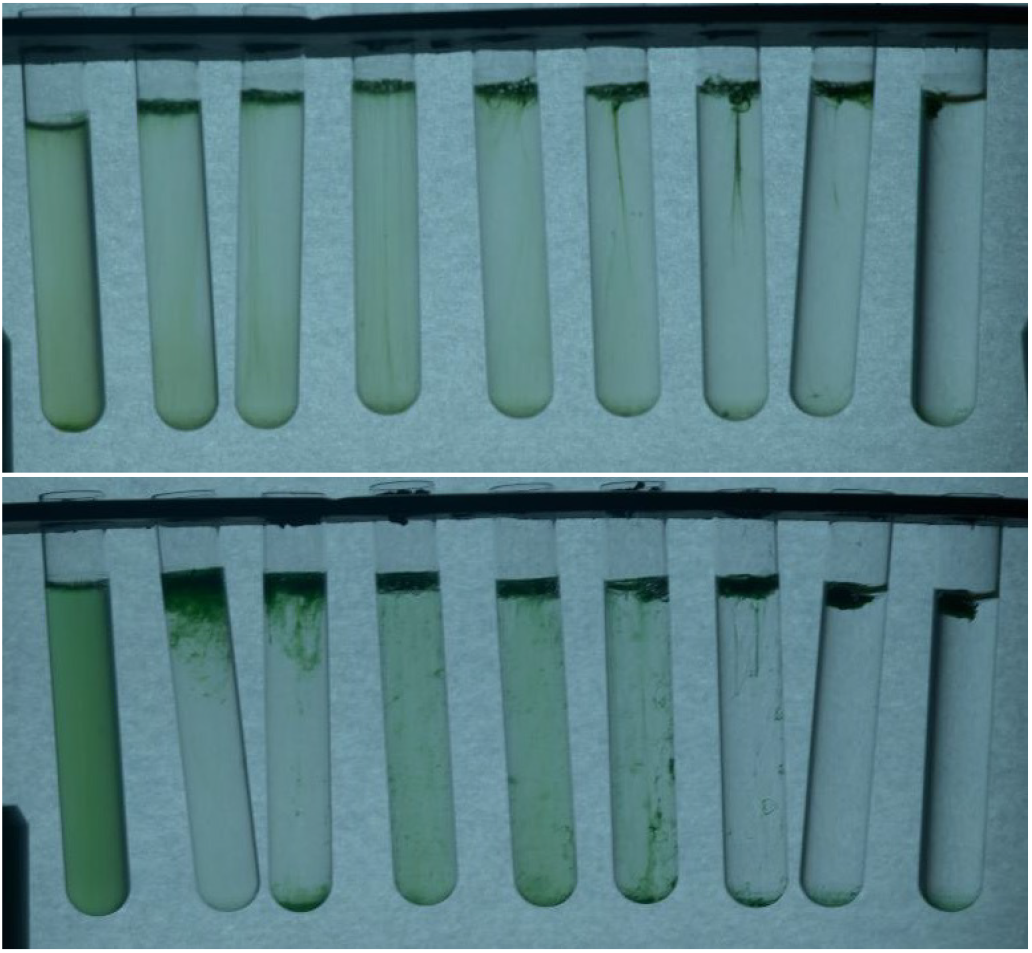
Samples of cultures of PCC 6803 in buffered media, initial OD = 1.3 and their evolution after addition of CaCl_2_ at various concentration. *Up* MOPS buffer 0 mM (control tube), 50 mM, 60 mM, 70 mM, 80 mM, 90 mM, 100 mM, 120 mM, 150 mM. *Down* CAPS buffer 0 mM (control tube), 0.5 mM, 1 mM, 1.5 mM, 2 mM, 2.5 mM, 3 mM, 5 mM and 10 mM.

#### Anabaena sp. PCC 7120

Intuitively, the filamentous shape of PCC 7120 can be though being an asset to exhibit IRMB, but the observations in unbuffered and buffered media did not meet these expectations, with some exception under very specific conditions. In unbuffered medium, the biomass is observed to sink relatively quickly (CS regime), whatever the added concentration in BG11 from 2 to 10X, see video in S.I. We repeat the attempt with OD_0_ significantly higher, at 1.72 instead of 0.6 keeping in mind that this would possibly strengthen the biomass cluster, which did not change much the observed regime. In rare cases, a tiny amount of biomass is stuck in bubbles at the surface, but the size of the cluster is far smaller than that for usual IRMB.

In buffered media, the observations are almost similar to those in unbuffered medium (CS). In CAPS and TAPS buffers with added BG11 at 5X concentration, similar tiny clusters remain stuck to the surface with bubbles, but most of the biomass sinks.

A remarkable exception, is when OD_0_ is taken significantly higher (around 2 or more) with a light intensity around 3000 Lux. In this case, the addition of CaCl_2_ induces a mixed behavior with CS below a threshold of 20 mM in added CaCl_2_, and IRMB+PS above this threshold. Let us note that even in CS, some quick rise-and-fall motion is observed, which testifies the biomass cluster are then not strong enough to sustain rising oxygen bubbles, see video in S.I. Therefore, CaCl_2_ is found to induce a stronger EPS matrix for the biomass.

### Synechococcus Elongatus PCC 7942

In unbuffered medium, PCC 7942 can react to the addition of BG11 with a specific regime : the intermittent rise and sink (IRS) of a significant part of the biomass. This regime occurs from 6X to 10X, and appears earlier at higher BG11 concentration. For concentrations of 5X or lower, the cultures remain homogeneous. Though the time-lapse videos do not give precise details on the mechanism, we assume this dynamics is caused by that the EPS matrix is strong enough to keep bubbles to grow inside by photosynthesis, but once bubbles get large enough the clusters reach the surface causing the bubbles to pop and the clusters then sink. A video of this regime is provided in S.I.. Operating with higher OD_0_ or higher light intensity does not qualitatively change the observations. Even at OD_0_ = 2.5 and 3000 Lux., the biomass just sink partially (PS) at BG11 higher than 6X, the other tubes remaining homogeneous for the duration of the experiment (1 day).

In buffered media, the cultures remain quasihomogeneous for most cases after addition of BG11, with only one exception in CAPS, where above 5X the culture shows almost complete sedimentation (CS) with only tiny clusters floating.

## CONCLUSIONS

For the four strains investigated here, the effect of pH on the growth of cyanobacterial cultures has revealed significant trends which, to the best of our knowledge, were up to now unreported.

First, the growth of any culture in unbuffered medium shows a clear common exponential growth during roughly the first 400 to 500 hours (no lag noticed), followed by a saturation phase when the OD can be up to 6. During this exponential phase, all cultures shows an increase of pH, from initial value around 6.5 to a final asymptotic value ranging between 10 and 11, and in some cases the pH value was measured up to 11.6 in a transient stage of the exponential phase. Depending on the initial OD, the pH is measured to grow constantly (OD_0_ *<* 0.1), to grow, reach a plateau value and slightly decrease (0.2 *<* OD_0_ *<* 0.1), or to grow and reach a plateau value (0.5 *<* OD_0_ *<* 0.6). The value of OD_0_ also influences the early stage growth of pH, as more concentrated cultures show a faster alcalinisation. The daily variations of pH show that the maximal value is reached at the end of the photosynthetic phase, while the respiration phase provokes the re-acidification of the cultures. The increase of pH could be attributed to the CCM which exports OH^−^ ions outside the cells. This hypothesis would require rigorous validation with in-situ measurements in the intracellular region. In environmental conditions with significant eutrophication, such an increase of pH could provide a competitive advantage to cyanobacteria and other photosynthetic microorganisms achieving CCM.

In buffered media, the cultures of all the four tested strains grow only in TAPS and CAPS, hence for pH higher than 9. In MES and MOPS buffers, the cultures barely survive at best, and often collapse. When measurable, the exponential phase appears less neatly and shorter than in unbuffered media, with sometime an early lag. The shortest doubling time are measured in CAPS, though their values are generally slightly higher than in unbuffered media for comparable OD_0_. This confirms the role of pH evidenced in unbuffered media, on which the culture itself creates the most advantageous conditions for its growth. Though, the constraint of a constant pH even in the respiration phase, could slightly limit the growth.

The influence of a buffered medium on the appearance of biomass aggregation and rising after salt addition, does not show any clear trend. For two investigated strains (PCC7005 and PCC6803), the different buffered media induce an indisputable influence, but the value of pH does not seem to correlate clearly with the observed regimes. For the two other strains (PCC7120 and PCC7942), the difference of observations between unbuffered and the different buffered media is marginal. Instead of a direct influence of the pH value, the molecules composing the buffer media could themselves induce a modification of the bloom-forming ionic bonding between cations and EPS. The chemical formula of all buffers are R-SO^−^_3_, and the sulfonate group is susceptible to form ionic bonds with Na^+^ or Ca^2+^, and consequently can modify or hinder the chelation of EPS with cations. The evidence of sulfated EPS in PCC6803 and their importance in bloom formation [37] also advocate for a non-neutral role of the buffers. Future studies should investigate on the influence of external environmental parameters on this chelation.

The cultures of PCC 7005, PCC 6803 and PCC 7120 originate from strains provided by Pr. A. Mejean (laboratoire LIED, University of Paris) and the cultures of PCC 7942 originate from strains provided by Pr. F. Chauvat (CEA Saclay).

PB conducted the experiments, PB and JD analysed the data, PB and JD wrote the manuscript.

## Notes

### Competing Interest Statement

The authors have declared no competing interest.

